# Short-term changes in a chicken comb vascular model mimicking port-wine stain after synchronous ultrasound parallel beam treatment: an exploratory study

**DOI:** 10.64898/2026.11.01.763470

**Authors:** Tingwei Zhang, Ruicai Wang, Zhengqiang Yang, Guosheng Wang

## Abstract

Synchronous ultrasound parallel beam (SUPERB) technology heats the mid-dermis without relying on hemoglobin as an optical chromophore, but its effect on the cutaneous microvasculature has not been reported. We examined short-term responses of a hypervascular chicken comb model to SUPERB treatment combined with applicator compression. In six adult roosters, two sites on the left comb surface each received a single pulse of 3 to 5 J after compression blanching, and the right surface served as a within-animal control. Full-thickness biopsies were taken immediately after treatment and on days 3 and 7; one rooster died on day 2, leaving five for analysis. Hematoxylin and eosin sections were assessed descriptively by a blinded pathologist, and vascular endothelial growth factor A (VEGF-A) positive area and Masson collagen area were quantified on day 7 at the animal level. Reduced microvascular caliber and fewer intraluminal erythrocytes were recorded at 1 of 10 sites immediately after treatment, 8 of 10 sites on day 3 and 8 of 9 evaluable sites on day 7. On day 7, VEGF-A positive area was lower on the treated side in all five animals (11.5% vs. 20.5%; mean paired difference −9.1 percentage points, 95% confidence interval −13.3 to −4.9; exact Wilcoxon p = 0.0625, the smallest value attainable with five pairs). Collagen area did not change consistently. Tissue temperature and perfusion were not measured, and no sham-compression group was included. These hypothesis-generating findings support controlled studies that separate the effects of compression and ultrasound energy.

## Introduction

Port-wine stain (PWS) is a capillary malformation of the skin. Somatic activating mutations in *GNAQ* have been identified in affected tissue [11]. Pulsed dye laser treatment relies on selective photothermolysis [1], but long-term follow-up shows that some lesions redarken after initial improvement [5]. Hypertrophic, recurrent and poorly responsive lesions therefore still call for energy delivery approaches that do not depend on the same mechanism.

Synchronous ultrasound parallel beam (SUPERB) technology (Sofwave) is used to treat facial lines, wrinkles and skin laxity [4, 13]. The device emits high-frequency, non-focused ultrasound from an array of parallel transducers and creates parallel thermal zones in the mid- dermis at a depth of approximately 1.5 mm, while contact cooling protects the epidermis [4]. In its 510(k) submission to the US Food and Drug Administration (FDA), the manufacturer reported a treatment depth of 1 to 2 mm and a focal tissue temperature of 60 to 70 °C [12]. Ultrasound energy deposition depends on the acoustic properties of tissue and does not require hemoglobin as an optical chromophore. Chromophore-independent approaches such as fractional CO_2_ laser have been used for hypertrophic PWS [14], and a computational study has examined the feasibility of photoacoustic-guided ultrasound treatment for PWS [2]. Direct experimental data on how SUPERB affects the cutaneous microvasculature are lacking.

The chicken comb is highly vascular, easy to sample and allows paired observation, and it has been used as a model in PWS-related photodynamic therapy research [7]. To our knowledge, the in vivo effect of SUPERB on the cutaneous microvasculature has not been reported. We used the chicken comb as a hypervascular skin model and examined short-term histology, vascular endothelial growth factor A (VEGF-A) immunoreactivity and collagen area after a single Sofwave treatment. Our aim was to determine whether a microvascular tissue response worth further testing could be detected.

## Materials and methods

### Study design and animals

This was an exploratory, non-randomized study with a within-animal control. Six adult farm-raised domestic roosters (approximately 24 months old; body weight approximately 2 kg) were enrolled. The experiment was conducted in the animal laboratory of Guangdong Laidi Biomedical Research Institute Co., Ltd., China. No formal sample size calculation was performed; the number of animals was based on feasibility and the principle of reducing animal use. In every animal, the left surface of the comb was treated, and the right surface received no ultrasound energy and served as the control. The left side was used in all animals so that treated sites could be identified reliably at follow-up; side allocation was not randomized. One rooster died on day 2 after treatment. No necropsy was performed, the cause of death was not determined, and its day 0 specimens were excluded from analysis. The remaining five animals were coded G, O, R, W and Y. Two treatment sites (coded 1 and 2) were placed on the left surface of each surviving animal, giving 10 sites in total. After the day 7 biopsies, the animals were euthanized according to the approved protocol. The protocol was approved by the Laboratory Animal Care and Ethics Committee of Guangdong Laidi Biomedical Research Institute Co., Ltd. (No. LDSW2025037-2; May 29, 2025), and the study is reported in accordance with the Animal Research: Reporting of In Vivo Experiments (ARRIVE) 2.0 guidelines [9].

### Treatment and tissue sampling

Local anesthesia was achieved with 1% lidocaine without epinephrine (10 mg/mL), injected at several points around the base of the comb at 0.1 to 0.2 mL per side. Ultrasound coupling gel was applied 1 to 2 minutes later. Treatment was delivered with the Sofwave system (Sofwave Medical Ltd., Israel). According to the operator manual, energy is set in joules per pulse over an adjustable range of 3 to 5 J. Each treatment site received a single pulse with an energy delivery time of 5 s. Energy settings for each site are listed in Table 1: both sites in animals G, R and W received 5 J, both sites in animal O received 4 J, and in animal Y site 1 received 3 J and site 2 received 4 J. Energy therefore varied between sites, and the study was not designed to compare energy levels. The FDA-cleared indication for Sofwave is aesthetic treatment of facial lines and wrinkles; its use on a vascular target in this study was investigational.

**Table 1.** Energy settings, sites quantified on day 7 and paired differences by animal.

| **Animal** | **Energy, site 1 / site 2** | **Sites quantified on day 7** | **VEGF-A difference (pp)** | **Masson difference (pp)** |
|---|---|---|---|---|
| G<sup>a</sup> | 5 J / 5 J | G1, G2 | −3.84 | −3.27 |
| O | 4 J / 4 J | O1, O2 | −12.06 | 3.22 |
| R | 5 J / 5 J | R1, R2 | −7.54 | 0.27 |
| W | 5 J / 5 J | W1 | −11.05 | 2.20 |
| Y | 3 J / 4 J | Y1 | −10.82 | 1.30 |

Differences are treated minus control. A sixth rooster died on day 2 and was not analyzed. No sections suitable for day 7 quantification were available for sites W2 and Y2. Multiple fields and regions of interest increased tissue sampling but did not increase the number of independent animals. VEGF-A, vascular endothelial growth factor A; pp, percentage points
<sup>a</sup>Site G1 was non-evaluable on day 7 hematoxylin and eosin staining; its VEGF-A and Masson values were included in the main analysis, and results without G1 are reported as a sensitivity analysis

Before energy delivery, the applicator was pressed onto the treatment area until visible blanching appeared (Fig. 1). The aim was to displace blood and reduce local perfusion temporarily, thereby reducing convective heat loss through blood flow so that ultrasound-deposited heat could accumulate in the tissue. Thermal models of ultrasound exposure show that blood flow can markedly alter tissue temperature distribution and the extent of thermal lesions [6]. This was a design assumption; tissue temperature and perfusion were not measured. The blanched area returned to red approximately 10 to 20 s after energy delivery ended and the applicator was removed. No purpura, edema or epidermal breakdown was observed immediately after treatment.

**Fig. 1.**
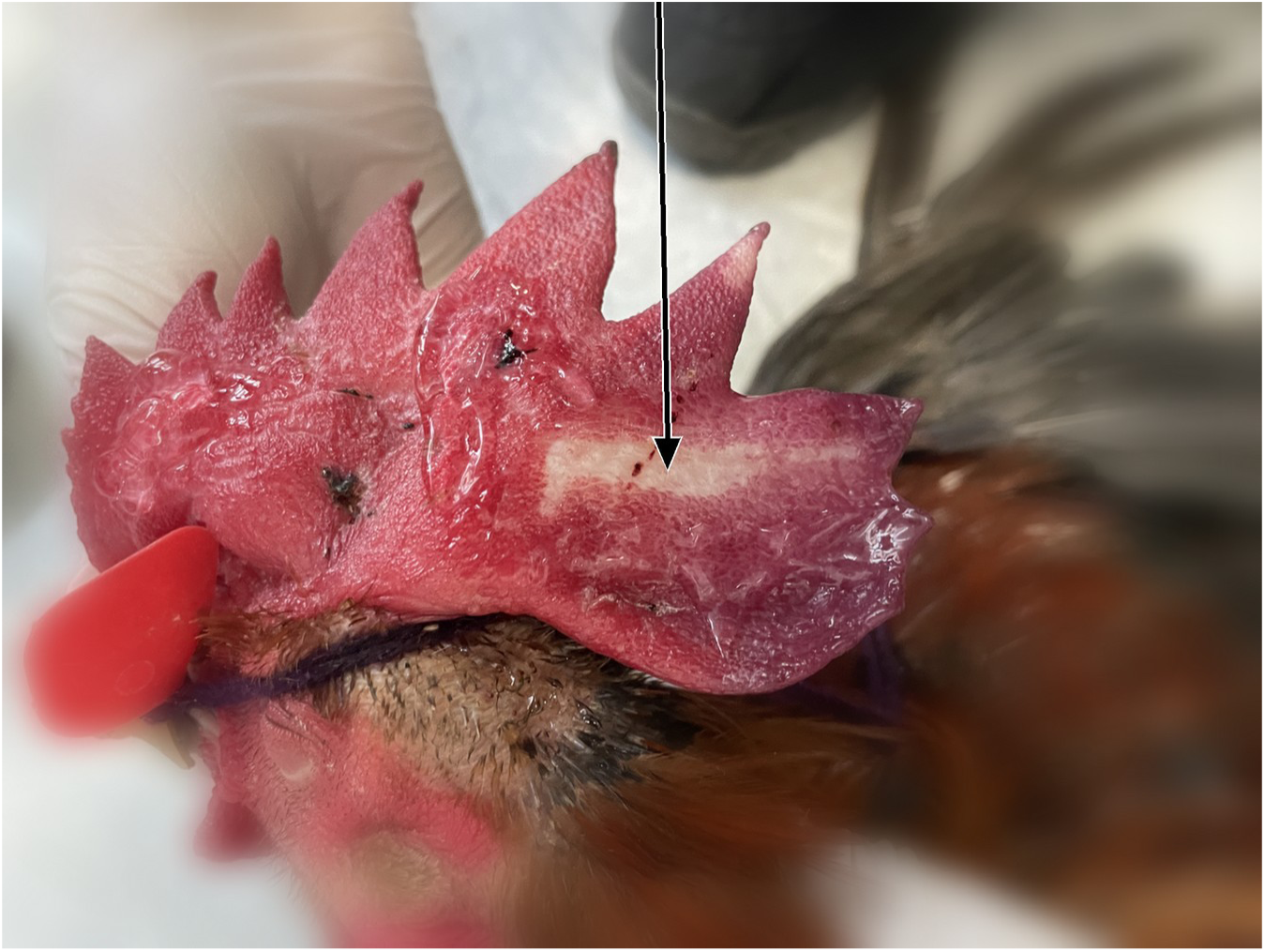
Compression-associated blanching during treatment. The arrow indicates visible blanching under applicator compression before energy delivery. This sign was used to confirm local blood displacement and is not evidence of vessel coagulation or of the extent of thermal injury. The background outside the comb was blurred; the comb was not edited

Full-thickness specimens were taken from each treatment site with a 10-mm biopsy punch immediately after treatment (day 0) and on days 3 and 7. Biopsies at different time points avoided previous biopsy wounds, with at least 3 mm between adjacent biopsy sites. Comb thickness at the sampling sites was approximately 1.5 to 2 cm. Each full-thickness specimen contained both the treated surface and the contralateral untreated surface, and control values were obtained from the opposite surface of the same section. Specimens were fixed in 10% neutral buffered formalin, embedded in paraffin and cut into 4-µm sections for hematoxylin and eosin (H&E) staining, Masson trichrome staining and VEGF-A immunohistochemistry. A specimen could contain several coded subsamples; for example, Y1-7-1 denotes animal Y, site 1, day 7, subsample 1. Sections labeled “BU” were replacement sections prepared after the original section failed quality control.

### Histopathological assessment

H&E slides were coded by a third person, and the pathologist was blinded to animal, time point and specimen orientation markings. One pathologist (R.W.) assessed reduction in microvascular caliber, reduction in intraluminal erythrocytes, collagen changes, inflammatory cells, differences between the two surfaces and endothelial proliferation. Percentages in the pathology records were visual estimates and were not used as morphometric measurements.

H&E findings were summarized descriptively, without composite scores or hypothesis testing. Items left blank were interpreted as “no obvious change,” as specified in the pathologist’s written notes. The day 7 specimen from site G1 was classified as non-evaluable because changes were present on both surfaces and the paired sides could not be identified reliably.

Fig. 2 shows representative day 7 H&E images from specimen Y1-7-1 (animal Y, site 1, 3 J). In the low-magnification overview, the untreated control surface is at the upper right and the Sofwave-treated surface is at the lower left. Original microscope calibration data could not be recovered, so these images are used for qualitative illustration only.

**Fig. 2.**
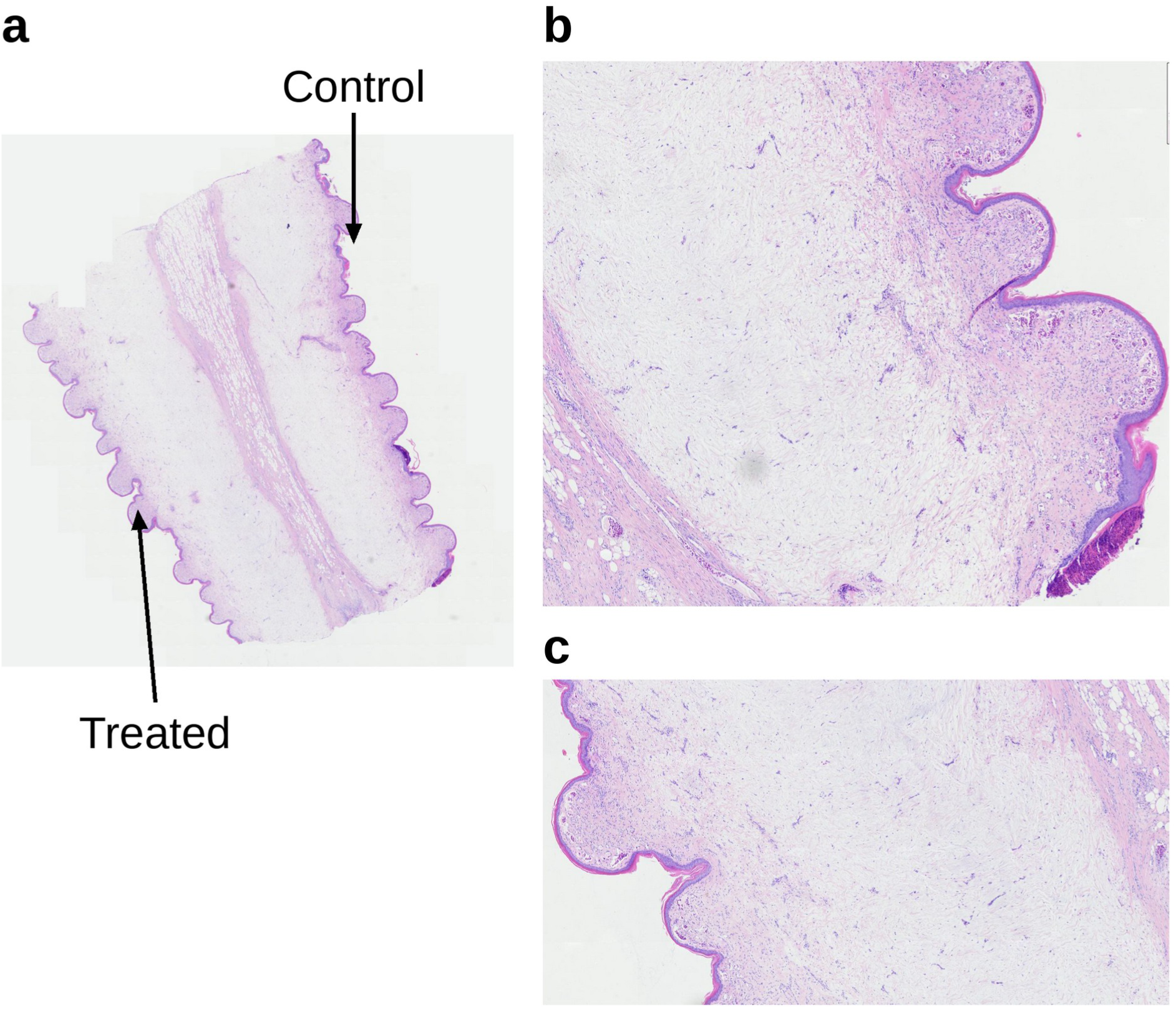
Representative day 7 hematoxylin and eosin images of specimen Y1-7-1 (animal Y, site 1, 3 J). **a** Full-thickness overview; the untreated control surface is at the upper right and the Sofwave-treated surface at the lower left. **b** Control-side detail. **c** Treated-side detail. All fields are from the same section. Original microscope calibration data could not be recovered, so no scale bar is shown and no absolute morphometry is implied. Panel c was reduced proportionally to the width of panel b

### VEGF-A immunohistochemistry

VEGF-A immunohistochemistry was performed by Tianjin Guosheng Zhongyuan Technology Co., Ltd. (Tianjin, China). Paraffin sections were dewaxed in xylene and rehydrated through graded ethanol. Antigen retrieval was performed in ethylenediaminetetraacetic acid (EDTA) buffer (pH 9.0) in a pressure cooker for 3 min, timed from the onset of steam release, followed by natural cooling. After three 3-min washes in phosphate-buffered saline (PBS, pH 7.4), endogenous peroxidase was blocked for 15 min at room temperature in the dark, and sections were blocked with 3% bovine serum albumin (BSA) for 1 h at room temperature.

Sections were incubated overnight at 4 °C in a humid chamber with a rabbit polyclonal anti- VEGF-A antibody (Bioss, bs-1313R; 1:300 in 3% BSA). After three 3-min washes in PBS, a goat anti-rabbit IgG secondary antibody (Servicebio, G1215-3; 1:200) was applied for 60 min at 37 °C. After further washes, freshly prepared 3,3′-diaminobenzidine (DAB) was applied with color development monitored under the microscope, and the reaction was stopped with tap water.

Nuclei were counterstained with hematoxylin for approximately 5 s, and sections were dehydrated through graded ethanol, cleared in xylene and mounted with neutral resin. VEGF-A immunoreactivity was read as brown DAB signal against blue nuclear counterstain. Slides were digitized with a Pannoramic Scan slide scanner (3DHISTECH). The laboratory report did not record negative (primary antibody omitted) or positive control sections. The manufacturer lists chicken as a predicted reactivity for this antibody, and chicken-specific validation was not performed in this study.

### Quantitative image analysis

On day 7, VEGF-A positive area fraction and Masson collagen positive area fraction were measured in ImageJ [10] by an observer blinded to side. Several regions of interest (ROIs) were placed in each micrograph, and VEGF-A ROIs avoided the epidermis and clearly non-vascular signal. Predefined thresholds were used (VEGF-A, 65 to 160; Masson, 0 to 120). Archived records showed that 10 of 725 VEGF-A ROI records and 70 of 600 Masson ROI records had been measured with other thresholds; these records were excluded in a sensitivity analysis. To avoid treating fields or ROIs from the same animal as independent biological replicates, ROI values were averaged within each image and then aggregated at the section, site and animal levels to give one control value and one treated value per animal. Repeated measurements of the same section were averaged first. Day 7 quantification covered eight sites (G1, G2, O1, O2, R1, R2, W1 and Y1); no sections suitable for quantification were available for sites W2 and Y2.

### Statistical analysis

Continuous variables are presented as mean ± standard deviation. The animal was the unit of analysis (n = 5). The primary effect estimate was the paired difference (treated minus control) with a 95% confidence interval (CI) based on the t distribution. Paired comparisons were also tested with the two-sided exact Wilcoxon signed-rank test. With five pairs, the smallest attainable two-sided p value for this test is 0.0625, so p < 0.05 cannot be reached regardless of effect size, and p values are reported as supporting information only. Two post hoc sensitivity analyses were performed: exclusion of site G1, whose side assignment was uncertain on H&E, and restriction to ROIs measured with the predefined thresholds. H&E findings were summarized descriptively. Analyses were performed in Python 3.13.5 with NumPy 2.3.5 and SciPy 1.17.0.

Preliminary results of this study were published as a conference abstract [15] that used an ROI-level analysis (treated 11.16 ± 6.93% vs. control 19.54 ± 8.15%, n = 21 ROI pairs, p < 0.0001). The present analysis uses the animal as the unit of replication, which accounts for the difference in the reported values.

### Use of large language models

ChatGPT (OpenAI; GPT-6, medium setting) and Claude (Anthropic; claude-sonnet-4-6 and claude-opus-5-5) were used to assist with literature searching, tabulation of archived ImageJ measurements, drafting of analysis and plotting scripts, figure panel layout and English language editing of author-written text. T.Z. reviewed and ran all scripts, and all reported values were calculated from the archived measurements. No AI tool generated experimental data, observations or image content. The authors checked all AI-assisted output and take full responsibility for the content of the manuscript.

## Results

### Animal outcomes and H&E findings

Of the six roosters enrolled, five were included in the histological analysis and one died on day 2. In the surviving animals, no persistent color change, purpura, edema or ulceration of the comb was observed on gross inspection immediately after treatment or on days 3 and 7. Reduced microvascular caliber and fewer intraluminal erythrocytes were recorded at 1 of 10 sites on day 0, 8 of 10 sites on day 3 and 8 of 9 evaluable sites on day 7 (Table 2). Site G1 on day 7 could not be interpreted as a reliable pair because both surfaces showed changes. Inflammatory cells, endothelial proliferation and collagen changes of varying degree were also recorded. Representative day 7 H&E findings from specimen Y1-7-1 are shown in Fig. 2.

**Table 2.** Descriptive hematoxylin and eosin findings by time point.

| **Time point** | **Evaluable sites** | **Reduced vessel caliber** | **Fewer intraluminal erythrocytes** | **Non-evaluable sites** |
|---|---|---|---|---|
| Day 0 (immediately after treatment) | 10 | 1/10 | 1/10 | 0 |
| Day 3 | 10 | 8/10 | 8/10 | 0 |
| Day 7 | 9 | 8/9 | 8/9 | 1 (G1)<sup>a</sup> |

A site was scored positive when the finding was clearly recorded in at least one coded histological subsample from that site. Sites were nested within animals, and no inferential testing of response rates was performed
<sup>a</sup>Both surfaces showed changes, and the paired sides could not be identified reliably

### VEGF-A immunohistochemistry

On day 7, mean VEGF-A positive area was 20.54 ± 1.78% on the control side and 11.48 ± 3.27% on the treated side, a mean paired difference of −9.06 percentage points (95% CI −13.26 to −4.86). The treated value was lower than the control value in all five animals (W = 0, p = 0.0625; Table 3, Fig. 3). After exclusion of site G1, all five animals still had lower treated values (mean difference −8.36 percentage points, p = 0.0625). Restricting the analysis to ROIs measured with the predefined thresholds gave a mean difference of −9.14 percentage points.

**Fig. 3.**
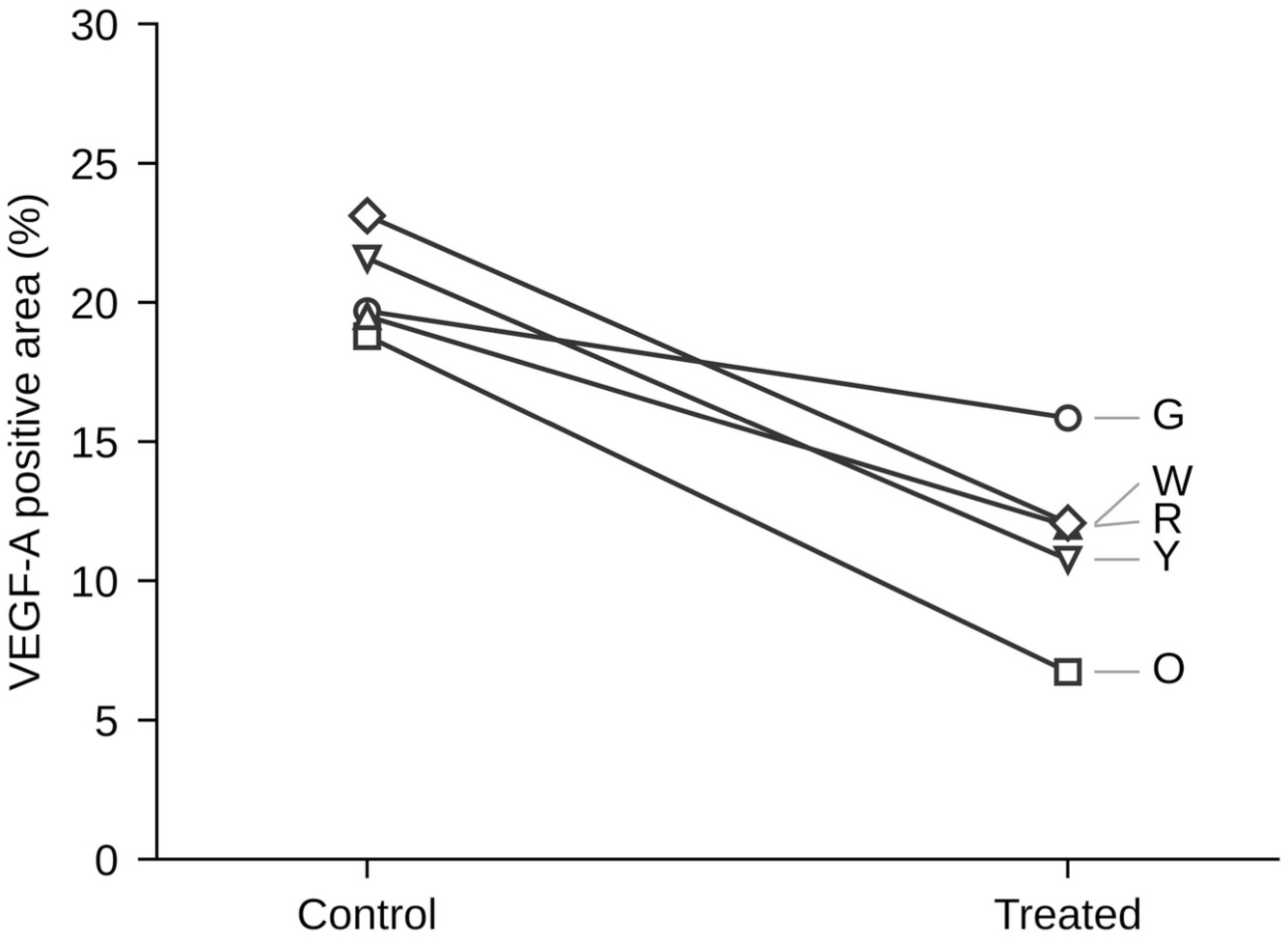
Animal-level paired vascular endothelial growth factor A (VEGF-A) positive area on day 7. Each line represents one animal (G, O, R, W and Y). The treated value was lower than the control value in all five animals; mean paired difference −9.06 percentage points (95% confidence interval −13.26 to −4.86); two-sided exact Wilcoxon signed-rank test, W = 0, p = 0.0625

**Table 3.**
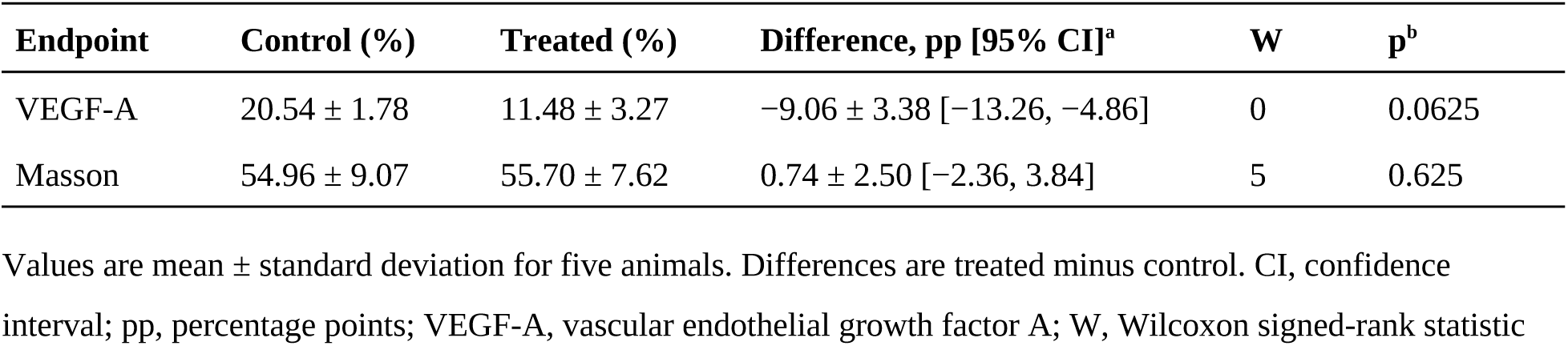
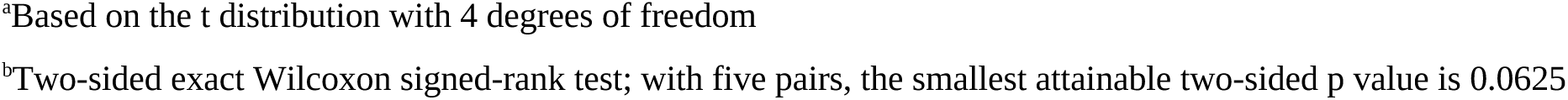
Animal-level quantitative results on day 7.

| **Endpoint** | **Control (%)** | **Treated (%)** | **Difference, pp [95% CI]<sup>a</sup>** | **W** | **p<sup>b</sup>** |
|---|---|---|---|---|---|
| VEGF-A | 20.54 ± 1.78 | 11.48 ± 3.27 | −9.06 ± 3.38 [−13.26, −4.86] | 0 | 0.0625 |
| Masson | 54.96 ± 9.07 | 55.70 ± 7.62 | 0.74 ± 2.50 [−2.36, 3.84] | 5 | 0.625 |

<sup>a</sup>Based on the t distribution with 4 degrees of freedom
<sup>b</sup>Two-sided exact Wilcoxon signed-rank test; with five pairs, the smallest attainable two-sided p value is 0.0625

Energy settings and paired differences for each animal are given in Table 1; the number of animals at each energy level was too small for comparison between energy levels. VEGF-A positive signal was located mainly in the vascular endothelium and perivascular tissue.

Values are mean ± standard deviation for five animals. Differences are treated minus control. CI, confidence interval; pp, percentage points; VEGF-A, vascular endothelial growth factor A; W, Wilcoxon signed-rank statistic

### Masson trichrome staining

On day 7, mean collagen positive area was 54.96 ± 9.07% on the control side and 55.70 ± 7.62% on the treated side, a mean paired difference of 0.74 percentage points (95% CI −2.36 to 3.84; W = 5, p = 0.625; Table 3, Fig. 4). The treated value was higher in four animals and lower in one. After exclusion of site G1, the treated value was slightly higher in all five animals (mean difference 2.56 percentage points, p = 0.0625), indicating that this endpoint was sensitive to the inclusion of a single site. Representative VEGF-A and Masson trichrome images from specimen Y1-7-1 are shown in Fig. 5.

**Fig. 4.**
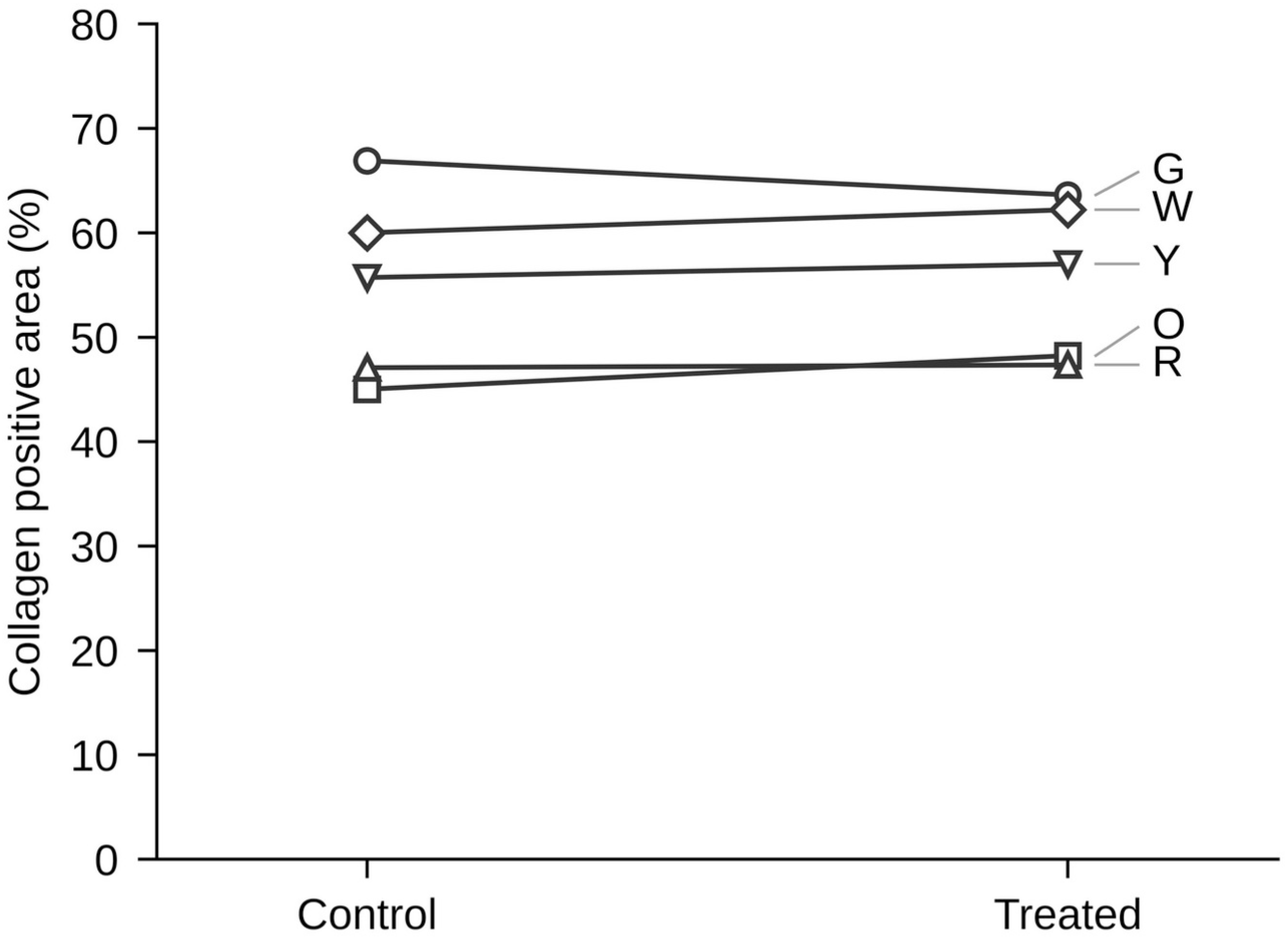
Animal-level paired collagen positive area (Masson trichrome) on day 7. Each line represents one animal. Mean paired difference 0.74 percentage points (95% confidence interval −2.36 to 3.84); two-sided exact Wilcoxon signed-rank test, W = 5, p = 0.625

**Fig. 5.**
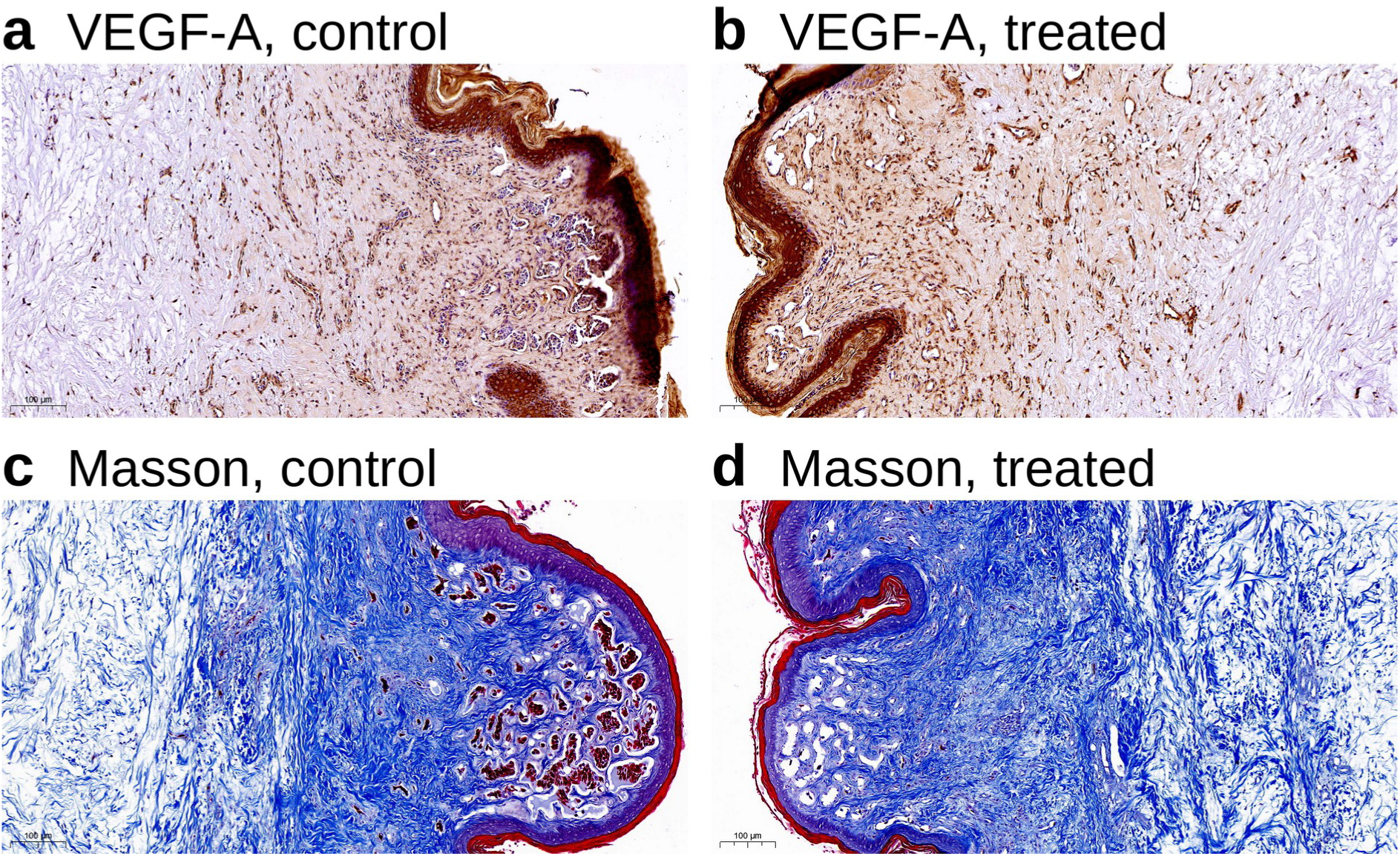
Representative VEGF-A immunohistochemistry and Masson trichrome images of specimen Y1-7-1 (animal Y, site 1, 3 J) on day 7. **a** VEGF-A, control side. **b** VEGF-A, treated side. **c** Masson trichrome, control side. **d** Masson trichrome, treated side. In a and b, brown indicates 3,3′-diaminobenzidine signal and nuclei are counterstained with hematoxylin. The original scanner scale bars (100 µm) are retained in each panel. Panels were placed and labeled outside the image area without local image adjustment. VEGF-A, vascular endothelial growth factor A

## Discussion

In this study, Sofwave treatment combined with applicator compression was followed by short-term microvascular changes in hypervascular chicken comb skin. VEGF-A positive area was lower on the treated side in all five surviving animals, and H&E records showed reduced microvascular caliber and fewer intraluminal erythrocytes at most evaluable sites on days 3 and 7. These changes were recorded at only 1 of 10 sites immediately after treatment, and their frequency rose on days 3 and 7. If the changes were mainly an acute mechanical artifact of compression, they would be expected to be most evident immediately after treatment; the increase over time fits an evolving tissue response better. Healing of earlier biopsy wounds in the same comb also evolves over time and may have influenced later sampling areas, so this confounder cannot be excluded. The sample size did not allow conventional significance to be reached in the animal-level two-sided test, but several endpoints pointed in the same direction. We propose that the main value of these results lies in a testable hypothesis: SUPERB combined with compression may affect the microvascular compartment of hypervascular skin.

Applicator compression was an integral part of the design. Its purpose was not to reduce “absorption” of ultrasound by hemoglobin, but to displace blood and lower perfusion temporarily so that less heat would be carried away by blood flow. Studies of ultrasound heating show that blood flow alters tissue temperature distribution and thermal dose and affects the extent of thermal lesions [6]. In a richly perfused tissue such as the chicken comb, compression may therefore favor local heat accumulation. This mechanism was the rationale for the design, and direct measurement of tissue temperature and perfusion during compression is a logical next step.

The present results do not allow the microvascular changes to be equated with vessel coagulation or clearance. A decrease in VEGF-A positive area may reflect a vascular biological response, but it does not show that vessels have been removed. In this study, VEGF-A signal was located mainly in the endothelium and perivascular tissue. VEGF-A is not an endothelium- specific marker, however, and keratinocytes, fibroblasts and inflammatory cells can also express it. The reactivity of the antibody with chicken tissue was predicted by the manufacturer and was not validated here, so the VEGF-A findings are treated as supporting evidence only. The reduced caliber and fewer intraluminal erythrocytes seen on H&E may also be influenced by edema, local tissue pressure, inflammation and short-term changes in blood flow.

From a thermal damage perspective, Doppegieter et al. found in isolated rat mesenteric arteries that 30 s of heating at 55.9 to 56.9 °C reduced endothelial, smooth muscle and vascular nerve function by half [3]. Thermal damage follows a time-temperature relationship: as temperature rises, the exposure needed for an equivalent damage integral shortens steeply, and Arrhenius-type models describe irreversible alteration of structural proteins above the hyperthermic range [8]. The manufacturer reports a focal tissue temperature of 60 to 70 °C [12], so an exposure of a few seconds could in theory match or exceed the vascular dysfunction produced by 30 s at 56 °C. Tissue temperature was not measured in this study, and the thermal damage kinetics of chicken comb vessels are unknown, so this inference remains a hypothesis. The study is therefore best read as exploratory evidence of a microvascular tissue response rather than proof of irreversible vascular injury.

Masson staining showed no consistent change in collagen area on day 7. This does not exclude changes in collagen structure or later remodeling, because area fraction does not capture fiber orientation, arrangement or maturation, and day 7 is an early time point. In the sensitivity analysis, exclusion of site G1 made the treated values slightly higher in all five animals, which shows that this endpoint depended heavily on a single site and should not be interpreted directionally at present. The more cautious reading is that no stable effect on collagen area was seen in the short term, whereas the microvascular measures showed a clearer directional change. Because the endpoints differ in nature, the contrast between the VEGF-A and Masson results cannot be taken as evidence that Sofwave acts selectively on vessels.

The chicken comb does not reproduce the genetic or developmental basis of human PWS, but its high vascular density makes it suitable for an initial look at how the cutaneous microvasculature responds to energy-based devices [7]. Our results support further mechanistic work. Future studies could include untreated, compression-only, Sofwave-only and Sofwave- plus-compression groups, use a uniform energy setting or a predefined energy gradient, and add validated chicken-reactive endothelial markers, objective measurement of vessel lumen area, perfusion or blood flow assessment, tissue temperature monitoring and longer follow-up. Only if such studies show that Sofwave consistently alters microvascular structure or perfusion after the effect of mechanical compression has been excluded would it be reasonable to discuss its translational potential in PWS.

### Limitations

This study has several limitations. The sample was small, with five animals analyzed; one further animal died on day 2 without necropsy, and the cause of death is unknown. The treated side was fixed on the left and not randomized, and no separate sham-compression control was included. Compression may have been transmitted through the comb, so the contributions of mechanical compression and ultrasound energy cannot be separated. Controls came from the opposite surface of the same comb. Because the comb was approximately 1.5 to 2 cm thick at the sampling sites and the thermal zone lies 1 to 2 mm below the treated surface, direct heat conduction to the control side is unlikely; however, transmitted pressure or diffusible factors acting on the control side would lead to underestimation of between-side differences. Energy varied between 3 and 5 J across sites, and sections for day 7 quantification were missing for sites W2 and Y2. The reactivity of the anti-VEGF-A antibody with chicken tissue was predicted by the manufacturer but not validated in this study, and negative and positive controls were not recorded. VEGF-A is not endothelium-specific, and validated endothelial markers, perfusion measurement and temperature monitoring were not used. Repeated biopsies of the same comb mean that wound healing may have influenced later time points. H&E was assessed descriptively by a single pathologist without assessment of inter-observer agreement, and the representative H&E images lack original scale calibration. A small number of ROI records were measured with thresholds other than the predefined values, although excluding them did not change the results. Comb appearance was assessed only by gross inspection, without colorimetric quantification.

Follow-up ended on day 7, so long-term vascular changes and collagen remodeling could not be assessed. These factors limit conclusions on mechanism, durability of vascular effects and clinical translation.

## Conclusion

In this exploratory study, Sofwave treatment combined with applicator compression was followed by short-term microvascular changes in hypervascular chicken comb skin. Reduced microvascular caliber and fewer intraluminal erythrocytes were recorded at most evaluable sites on days 3 and 7, VEGF-A positive area was lower on the treated side in all five animals, and collagen area did not change consistently on day 7. Because the antibody was not validated in chicken tissue, the VEGF-A results serve only as supporting evidence. These hypothesis- generating findings support mechanistic studies with sham-compression controls, uniform energy settings, blood flow and temperature monitoring and validated endothelial markers. The present evidence does not establish vessel clearance, durable vascular destruction or clinical efficacy in PWS.

## Statements and Declarations

### Competing

The authors have no relevant financial or non-financial interests to disclose. The Sofwave device was rented at the authors’ own expense, and the manufacturer and its distributors had no role in study design, data analysis or manuscript preparation.

### Ethics approval

All animal procedures were approved by the Laboratory Animal Care and Ethics Committee of Guangdong Laidi Biomedical Research Institute Co., Ltd. (No. LDSW2025037-2; May 29, 2025) and were performed in accordance with the approved protocol. Treatment was performed under local anesthesia, and the animals were euthanized after the day 7 biopsies according to the approved protocol. The study is reported according to the ARRIVE 2.0 guidelines.

### Consent to participate

Not applicable.

### Consent to publish

Not applicable.

### Data availability

The ImageJ measurement records and original section images that support the findings of this study are available from the corresponding author on reasonable request and will be deposited in a public repository upon journal publication.

### Code availability

The Python analysis and plotting scripts are available from the corresponding author on reasonable request and will be deposited together with the data upon journal publication.

### Author contributions

Conceptualization: Tingwei Zhang; Methodology: Tingwei Zhang; Investigation: Tingwei Zhang, Zhengqiang Yang (experiments and data collection), Ruicai Wang (histopathological assessment), Guosheng Wang (animal procedures and specimen preparation); Data curation: Tingwei Zhang; Formal analysis: Tingwei Zhang; Validation: Ruicai Wang; Visualization: Tingwei Zhang; Resources: Guosheng Wang; Project administration: Tingwei Zhang; Writing - original draft preparation: Tingwei Zhang; Writing - review and editing: Tingwei Zhang, Zhengqiang Yang, Ruicai Wang, Guosheng Wang. All authors read and approved the final manuscript.

## Acknowledgments

We thank Dr. Min Ding for assistance with immunohistochemical staining and Dr. Tingting Hu for assistance with histopathological image analysis. Preliminary results of this study were presented as an ePoster at the 2026 Annual Conference of the American Society for Laser Medicine and Surgery (ASLMS) and published in abstract form.

